# Specific protein-membrane interactions promote the export of metallo-β-lactamases via outer membrane vesicles

**DOI:** 10.1101/2021.03.18.436108

**Authors:** Carolina López, Alessio Prunotto, Guillermo Bahr, Robert A. Bonomo, Lisandro J. González, Matteo Dal Peraro, Alejandro J. Vila

## Abstract

Outer membrane vesicles (OMVs) act as carriers of resistance determinants such as metallo-β-lactamases. The metallo-β-lactamase NDM-1 is present in OMVs produced by Gram-negative bacteria since it is a lipidated, membrane-anchored protein. The soluble domain of NDM-1 also forms electrostatic interactions with the membrane. Herein, we show that these interactions promote its export into OMVs produced by *Escherichia coli*. We report that favorable electrostatic protein-membrane interactions are also at work in the soluble enzyme IMP-1, while being absent in VIM-2. These interactions correlate with an enhanced secretion into OMVs of IMP-1 compared to VIM-2. Disruption of these interactions in NDM-1 and IMP-1 impairs export into vesicles, confirming their role in defining the protein cargo in OMVs. These results also indicate that export of metallo-β-lactamases into vesicles in their active form is a common phenomenon that involves cargo selection based on molecular features.

## INTRODUCTION

The rise of carbapenem-resistant bacteria producing metallo-β-lactamases is of great concern since this class of β-lactam antibiotics is reserved as a “last-resort” option for life-threatening infections (1, 2). Metallo-β-lactamases (MBLs) represent the largest family of carbapenemases (3–6). Among MBLs, the plasmid-borne NDMs, VIMs and IMPs are the enzymes with the highest clinical relevance and geographical dissemination (3, 5, 7). This genetic localization has favored their dissemination into different opportunistic and pathogenic bacteria (8–10).

MBLs from the NDM family are lipoproteins anchored to the outer membrane (OM) of Gram-negative bacteria (11, 12). This cellular localization enables the transport of NDM-1 into outer membrane vesicles (OMVs) (12). OMVs are spherical lipid bilayer nanostructures released by all Gram-negative bacteria (13) which are recognized as “public goods”, as they can transport different molecules outside the boundaries of the bacterial cell and share them with neighboring communities of bacteria (14, 15). Ciofu and coworkers reported for the first time the secretion of β-lactamases into OMVs (16). Later studies in recent years have strengthened this observation, showing that OMVs from different bacteria are able to transport serine-β-lactamases, MBLs and genes coding for these enzymes (16–21). These observations suggest that this phenomenon is common rather than being an exception and supports the hypothesis that these vesicles play a pivotal role in the dissemination of bacterial antibiotic resistance. However, the features that govern the selection of the protein cargo into OMVs are largely unknown.

NDM-1 is exported via OMVs produced by different bacteria as a fully active enzyme, endowing the vesicles with a potent carbapenemase activity (12, 22). NDM-1-loaded vesicles are able to inactivate the antibiotic in the near environment and to protect nearby populations of bacteria susceptible to antibiotics (12). NDM-1 is selectively transported into vesicles by being anchored to the outer membrane (12). However, a soluble variant engineered in the lab (NDM-1 C26A, lacking the lipidated Cys residue) is also present in vesicles (although in smaller amounts) (12). This observation compelled us to ask whether there are other molecular features in addition to membrane anchoring that favor the cargo selection into vesicles. Indeed, the globular domain of NDM-1 was shown to interact with the negatively charged anionic lipids in the bacterial membrane, through electrostatic interactions mediated by two Arg residues: Arg45 and Arg52, which are conserved in all known NDM variants (23). Substitution of these two residues to negatively charged Glu disrupts the interaction with the lipid bilayer (23). Here, we show that these interactions are relevant in determining the amount of active NDM-1 present in *Escherichia coli* vesicles. We also analyzed the protein-membrane interactions of IMP-1 and VIM-2, clinically relevant MBLs that are soluble periplasmic proteins. We report that attractive electrostatic interactions between the membrane and the soluble domains of MBLs favor their export into OMVs, as is the case for NDM-1 and IMP-1. Instead, in VIM-2 the lack of interaction with the membrane results in an impaired export into vesicles. Disruption of the membrane interactions in NDM-1 and IMP-1 induce a drop in the amount of secreted MBL, confirming that these are key protein features active in selecting the protein cargo into vesicles.

## RESULTS AND DISCUSSION

### The NDM-1 cargo into OMVs is promoted by specific electrostatic interactions with the membrane

In order to study the contribution of specific interactions between the bacterial membrane and NDM-1 in its export via vesicles, we analyzed the amount of active protein in the OMVs produced by *E. coli* cells expressing: (1) the native, lipidated, membrane bound NDM-1, (2) a soluble variant in which the lipidation site is removed (NDM-1 C26A) and (3) a lipidated double mutant in which the two Arg residues (Arg45 and Arg52) are replaced by Glu (NDM-1 2RE). All studied MBLs were fused to a C-terminal Strep-tag sequence (-ST) for immunoblotting quantification. OMVs were purified from the supernatants of cell cultures of *E. coli* and then analyzed by SDS-PAGE and immunoblotting with anti-ST antibodies. Figures 1 B-C show that removal of the lipidation site in NDM-1 C26A leads to a 70% drop in the amount of transported NDM-1, in agreement with previous experiments (12). The NDM-1 2RE mutant, despite preserving the membrane localization, displays an impaired transport of almost 35% (Fig. 1B-C). Remarkably, the R45E/R52E mutation did not alter the ability of this enzyme to confer resistance against β-lactams in *E. coli* cells (Table S1), nor its carbapenemase activity within vesicles (Fig. 1D). Considering that these mutations disrupt the electrostatic interaction of NDM-1 with the membrane, we conclude that interactions between the globular domain of NDM-1 and the bacterial outer membrane enhance the secretion of this enzyme into vesicles. These interactions may act as a “fine-tuning” mechanism for cargo selection of lipoproteins.

**Figure 1.**
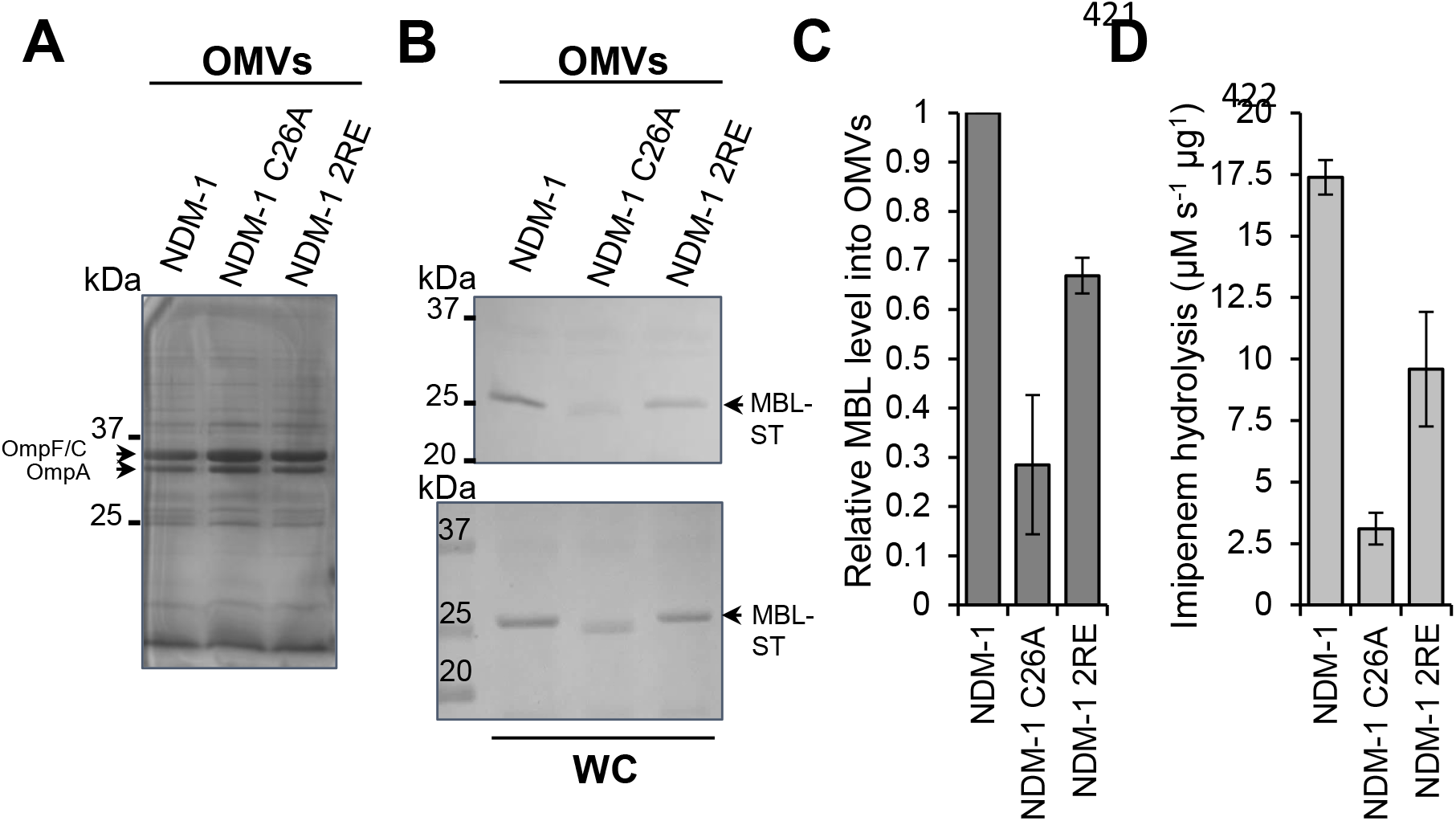
The specific interaction between NDM-1, mediated by residues Arg45 and Arg52, and the bacterial membrane promotes its export into vesicles. (A) SDS-PAGE of the OMVs purified from *E. coli* expressing *bla*_NDM-1_, *bla*_NDM-1 C26A_ or *bla*_NDM-1 2RE_ .(B) Immunoblotting detection of NDM-1, NDM-1 C26A and NDM-1 2RE (fused to a C-terminal Strep-tag sequence, -ST) in OMVs and in whole cells (WC) from *E. coli* strains expressing each MBL. (C) Protein levels of NDM-1, NDM-1 C26A and NDM-1 2RE into OMVs. The plotted values were calculated as described in the materials and methods section. Data correspond to three independent experiments and are shown as the mean value. Error bars represent the standard deviation (SD). (D) Imipenem hydrolysis rate by OMVs purified from *E. coli* expressing NDM-1, NDM-1 C26A or NDM-1 2RE. Data correspond to two independent experiments and are shown as the mean value. Error bars represent the standard deviation (SD).

### The MBL cargo into OMVs depends on the protein-membrane affinity

We hypothesized that electrostatic interactions of soluble periplasmic MBLs with the bacterial membrane may also favor transport of these proteins into OMVs, regardless of a lipid anchor. So, we analyzed the affinity of WT NDM-1 and soluble, periplasmic VIM-2 and IMP-1 with the bacterial membrane through coarse-grained (CG) molecular dynamics (MD) simulations (Fig. 2A). The membrane bilayer was modeled mimicking the lipid composition of the bacterial outer membrane, as previously reported (23). IMP-1 was predicted to have a strong interaction with the membrane, as evidenced by the same number of binding events as NDM-1 (Fig. S1A and Fig. 2A). In general, IMP-1 spent a large portion of simulation time in proximity of the bilayer (Fig. S1B). In contrast, VIM-2 did not show significant binding events in any of the MD replicas (Fig. S1A-B and Fig. 2A). In the case of IMP-1, the interaction occurred by means of a specific patch spanning from Gly85 to Ser94 and from Thr133 to Pro153. Although this region is different to that previously identified for NDM-1, it also features a cluster of positively charged residues, in particular four lysine residues (Lys87, Lys89, Lys145 and Lys147) (Fig. 2A).

**Figure 2.**
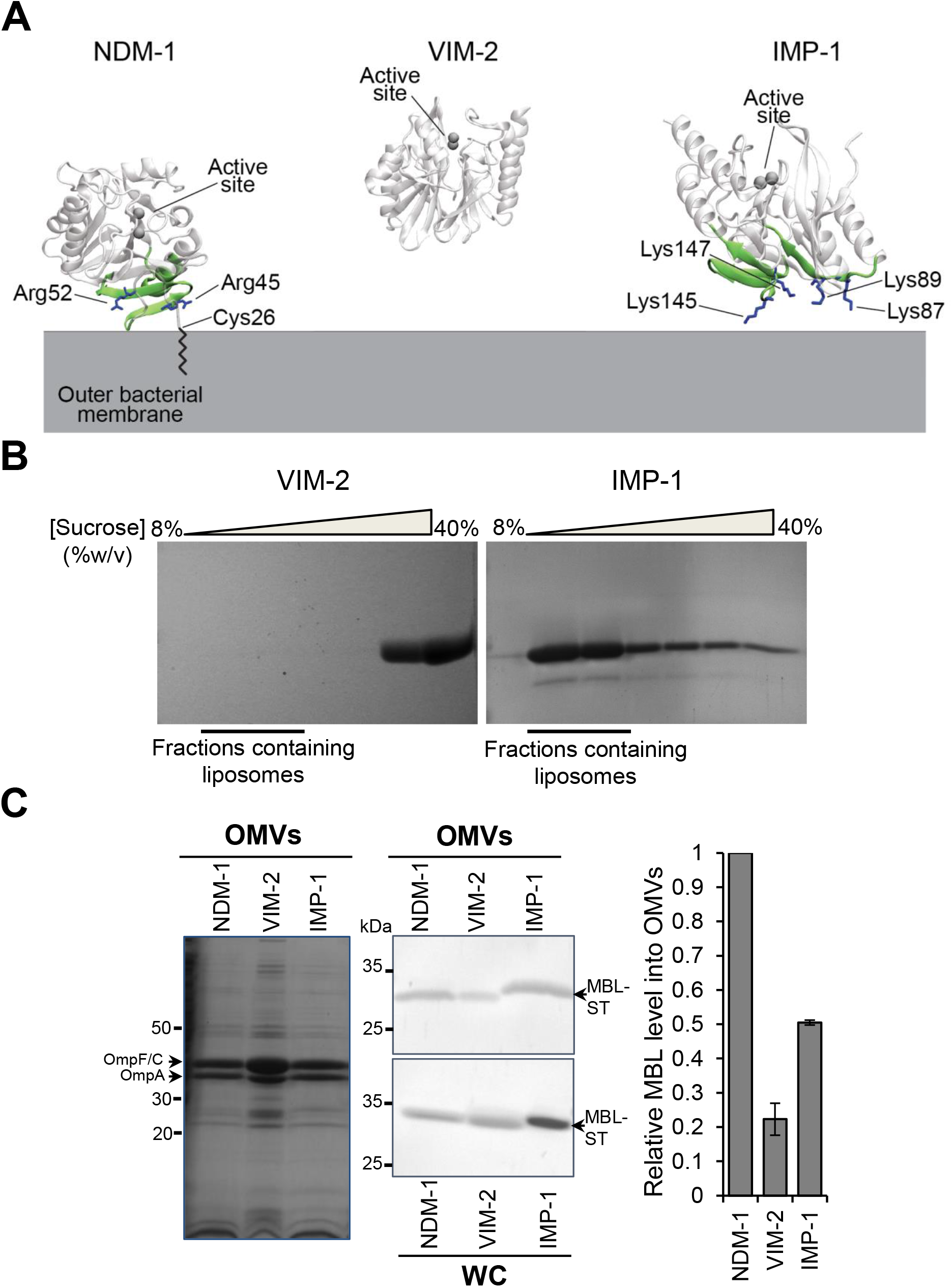
NDM-1, IMP-1 and VIM-2 MBLs have different affinities for the bacterial membrane that impact on export into vesicles. (A) Identified surfaces of interaction between NDM-1, VIM-2 and IMP-1 and the bacterial outer membrane. The domains that interact directly with the membrane are depicted in green cartoon, whereas the positively charged residues that drive the interaction are in blue sticks. VIM-2 did not interact with the membrane in any of the MD replicas, and therefore does not present an interaction patch. (B) SDS-PAGE analysis of sucrose gradient fractions from liposome flotation assays of VIM-2 and IMP-1. The flotation assays were carried out using liposomes made with an outer membrane composition from *E. coli*. (C) SDS-PAGE analysis (*Left panel*) and immunoblot analysis (*center panel*) of OMVs and whole cells (WC) from *E. coli* expressing NDM-1, VIM-2 or IMP-1, after induction with 20 µM IPTG. *Right panel*: comparison between levels of NDM-1, VIM-2 and IMP-1 into OMVs. The plotted values, relativized to the NDM-1 values, were obtained as described in the materials and methods section. Data correspond to two independent experiments and are shown as the mean value. Error bars represent the standard deviation (SD).

To experimentally test these predictions, we studied the interaction of IMP-1 with the membrane *in vitro* by means of liposome flotation assays (23), incubating soluble IMP-1 with liposomes mimicking the lipid composition of the bacterial outer membrane (Fig. 2B). Samples were loaded at the bottom of a sucrose gradient and were then ultracentrifuged. In these experiments, liposomes float towards lower concentrations of sucrose, and free proteins remain at the bottom of the gradient. SDS PAGE analysis of these samples showed that IMP-1 co-localizes with liposomes, i.e., IMP-1 is strongly associated with the membrane (Fig. 2B). VIM-2, instead, was largely found in the fractions corresponding to unbound proteins (Fig. 2B). Next we sought to evaluate if the interaction of IMP-1 with membranes favors its transport into vesicles. We compared the levels of IMP-1, NDM-1 and VIM-2 transported into vesicles from *E. coli* cells expressing these enzymes. Despite being a soluble protein, the amount of IMP-1 in OMVs is 50% of the transported levels of the membrane-bound NDM-1 (Fig. 2C). This is significant when compared to other soluble MBLs, such as VIM-2 (Fig. 2C) or the soluble variant NDM-1 C26A itself. The protein levels of VIM-2 and NDM-1 C26A exported into vesicles were less than half of those observed for IMP-1.

### The interaction of the Lys rich patch in IMP-1 contributes to its export via OMVs

The results obtained heretofore indicate that electrostatic interactions leading to membrane association are responsible for boosting the transport of MBLs into OMVs, regardless of the presence of a lipid anchor. To test this hypothesis, we decided to disrupt the interactions of IMP-1 with the membrane by replacing the 4 Lys residues (Lys87, Lys89, Lys145 and Lys147) by Glu residues. CG MD simulations of this IMP-1 mutant (IMP-1 4KE) predict a significant drop in the number of binding events compared to WT IMP-1 (3 out of 5 replicas did not show any binding, Fig. S1A), and in general a lower affinity towards the membrane, as highlighted by the significantly lower amount of simulation time spent by the enzyme in the proximity of the bilayer (Fig. S1B). Then, we evaluated the amount of IMP-1 and IMP-1 4KE mutant secreted into vesicles from *E. coli* cells expressing these MBLs. Figure 3A-C shows that the export via vesicles of IMP-1 4KE experiences a decay of almost 35% with respect to WT IMP-1, confirming the role of these specific electrostatic interactions in promoting the IMP-1 cargo into OMVs. Notably, the resistance profile (Table S1) and the carbapenemase activity (Fig. 3D) displayed by IMP-1 and the IMP-1 4KE mutant were preserved in *E. coli* cells and within the vesicles, indicating that these mutations did not alter the folding nor metal binding capability of the enzyme. Remarkably, the Lys residues at positions 89 and 147 are fully conserved among the 80 allelic variants in the IMP family (24). Lys145 is replaced by an Asn in only 2 allelic variants, while Lys87 is conserved in 65 variants (Fig. S2). This high degree of conservation predicts that members of the IMP family may be tuned to interact favorably with the bacterial membrane and consequently be exported through vesicles.

**Figure 3.**
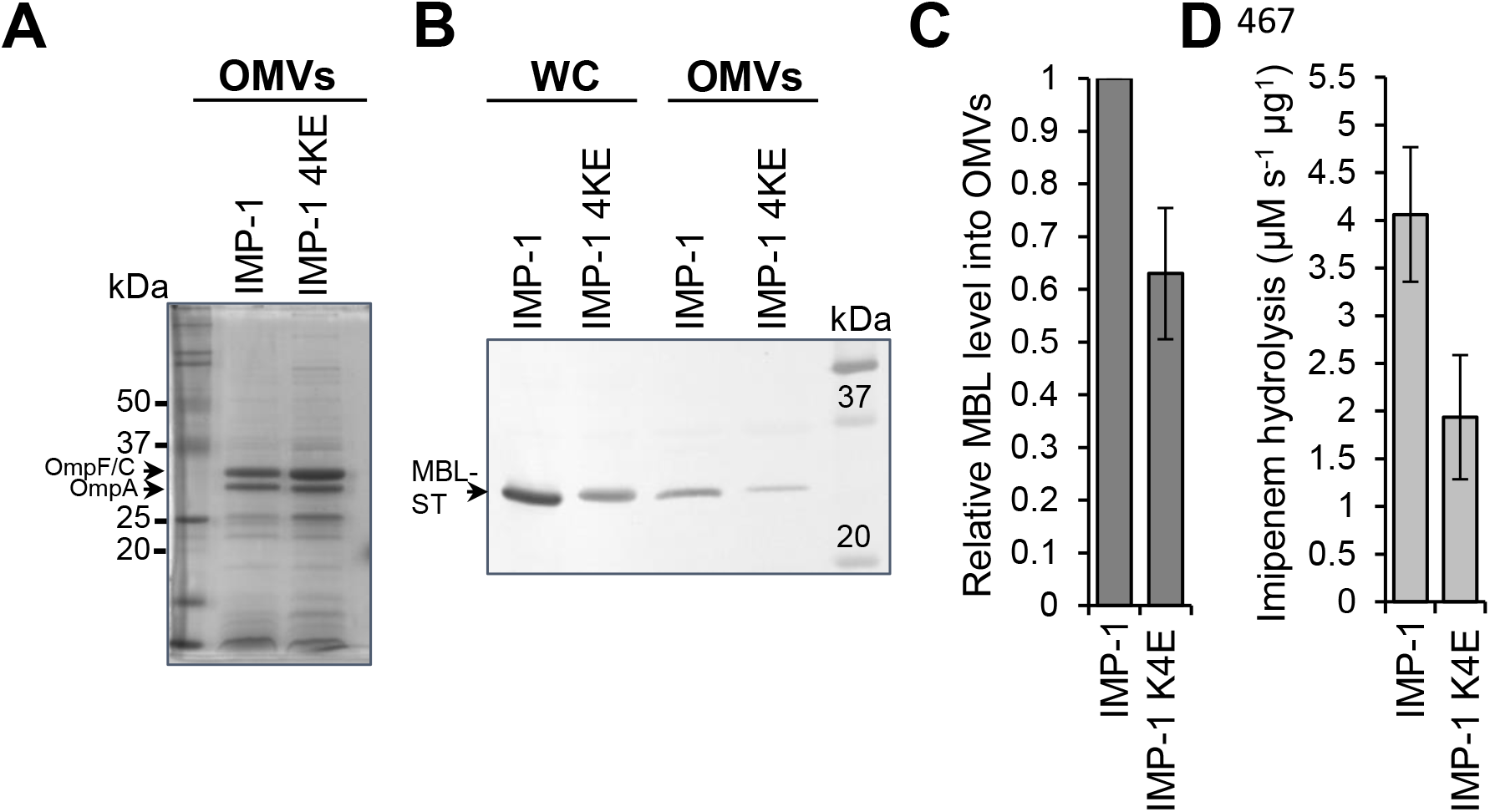
The specific interaction between IMP-1, mediated by Lys87, Lys89, Lys145 and Lys147, and the bacterial membrane promotes its export into vesicles. (A) SDS-PAGE analysis of OMVs from *E. coli* expressing IMP-1 and IMP-1 4KE mutant (IMP-1 K87E/K89E/K145E/K147E), (B) immunoblot analysis of OMVs and whole cells (WC) from *E. coli* expressing IMP-1 or IMP-1 4KE. (C) Comparison between levels of WT IMP-1 and IMP-1 4KE into OMVs. The plotted values, relativized to the IMP-1 value, were obtained as described in the materials and methods section. Data correspond to three independent experiments and are shown as the mean value. Error bars represent the standard deviation (SD). (D) Imipenem hydrolysis rate by OMVs purified from *E. coli* expressing IMP-1 or IMP-1 4KE. Data correspond to two independent experiments and are shown as the mean value. Error bars represent standard deviation (SD).

## CONCLUDING REMARKS

OMVs have been reported to transport MBLs in an active form, contributing to resistance by hydrolyzing antibiotics *in situ* and by protecting bacterial populations otherwise susceptible to antibiotics (12, 25). Elucidation of the molecular features that contribute to export of the active forms are key to understand the role of OMVs in antibiotic resistance. The current results reveal a direct correlation between attractive electrostatic interactions of MBLs with the bacterial outer membrane and the protein levels transported into OMVs in an active form. These features add to the already reported effect of membrane anchoring in NDM-1 in favoring export to vesicles and to recent findings in *Bacteroides* spp. reporting the impact of the efficient packaging of lipoproteins for OMVs sorting (26). Overall, this reveals that there are multiple molecular features active in synergy in the OMV cargo selection related to antimicrobial resistance that deserve further attention.

## MATERIALS AND METHODS

### Bacterial strains, culture conditions and construction of MBL mutants

*Escherichia coli* DH5α was used for expression of the pMBLe-Gm^R^ vector, as an empty vector and including the genes coding for the different MBLs: NDM-1, VIM-2 and IMP-1 and their mutants, NDM-1 C26A, NDM-1 R45E/R52E (NDM-1 2RE) and IMP-1 K87E/K89E/K145E/K147E (IMP-1 4KE). All variants were cloned in the full-length version into pMBLe, fused to a C-terminal Strep-tag II sequence and under the control of a β-IPTG-inducible pTac promoter, as previously described (12, 22). Cells were routinely grown aerobically at 37°C in lysogeny broth (LB) or on LB agar plates, except when indicated. Gentamicin was used when necessary at 20 µg/ml. The pMBLe-NDM-1 2RE-ST was obtained from plasmid pMBLe-NDM-1-ST through site-directed mutagenesis by plasmid amplification using primers NDM-1 2RE Fw (5’ GAAACTGGCGACCAAGAGTTTGGCGATCTGGTTTTCGAGCAGCTCGCACCG AATG 3’) and NDM-1 2RE Rv (5’ CATTCGGTGCGAGCTGCTCGAAAACCAGATCGCCAAACTCTTGGTCGCCAG TTTC 3). Plasmid pMBLe-IMP-1 K87EK89E-ST was obtained by directed mutagenesis by plasmid amplification on plasmid pMBLe-IMP-1-ST, using primers IMP-1 K87EK89E Fw (5’ GGTTTGTGGAGCGTGGCTATGAAATAGAAGGCAGCATTTCCTCTC 3’) and IMP-1 K87EK89E Rv (5’ GAGAGGAAATGCTGCCTTCTATTTCATAGCCACGCTCCACAAACC 3’). Then, using the latter plasmid as a template, the pMBLe-IMP-1 4KE-ST was obtained, using primers K145EK147E Fw (5’ GGAGTTAACTATTGGCTAGTTGAAAATGAAATTGAAGTTTTTTATCCAGG 3 ‘) and K145EK147E Rv (5’ CCTGGATAAAAAACTTCAATTTCATTTTCAACTAGCCAATAGTTAACTCC 3’).

### Liposome preparation

Pure lyophilized phospholipids (1-palmitoyl-2-oleoyl phosphatidylethanolamine, tetraoleoyl cardiolipin, and 1-palmitoyl-2-oleoyl-phosphatidylglycerol) were purchased from Avanti Polar Lipids. The lipids were dissolved in chloroform and after mixing the required proportions of each pure lipid, the lipid mixtures were dried under a nitrogen atmosphere and then kept under vacuum for 2h. The dried lipid films were hydrated with 50 mM HEPES pH 7 and heated at 65°C for 1h with periodic vortexing. Lipid suspensions were frozen in liquid nitrogen and then thawed at 65°C, for a total of 5 cycles, and afterwards were passed through a 400 nm polycarbonate filter using an Avanti Miniextruder apparatus (Avanti Polar Lipids) at 65°C, with >20 passes through the device.

### Liposome flotation assay

Samples containing 55 µM purified protein (VIM-2 or IMP-1) in 50mM HEPES at pH 7 were incubated with liposomes for 30 min at room temperature. Sucrose was added to 40% w/v, the samples were loaded in an ultracentrifuge tube, and sucrose 25% w/v and sucrose 8% w/v (both buffered with 50mM HEPES pH 7) were layered on top, forming a discontinuous sucrose gradient. Afterwards, samples were ultracentrifuged for 1h at 4°C and 125000 x *g* in a Beckman SW Ti90 rotor, and fractions along the gradient were analyzed by SDS-PAGE to assess the final distribution of the MBL protein.

### MIC determinations

Ceftazidime (CAZ) and imipenem (IMI) MIC determinations for *E. coli* DH5α cells expressing the different MBLs, their mutants or carrying the pMBLe empty vector (EV) lacking any MBL gene (as a control), were carried out in lysogeny broth (LB) following the broth microdilution method according to the Clinical and Laboratory Standards Institute (CLSI) protocols (27).

### Purification of OMVs and quantification of MBLs levels into OMVs

300 ml of LB medium were inoculated with 3 ml of saturated *E. coli* pMBLe-*bla* culture, grown at 37°C up to an OD600 of 0.4, induced with 20µM IPTG, and growth continued overnight with agitation. The cells were harvested, and the supernatant filtered through a 0.45-µm membrane (Millipore). Ammonium sulfate was added to the filtrate at a concentration of 55% w/v, followed by overnight incubation with stirring at 4°C. Precipitated material was separated by centrifugation at 12,800 x *g* for 10 min, resuspended in 10 mM HEPES, 200 mM NaCl at pH 7.4, and dialyzed overnight against > 100 volumes of the same buffer. Next, samples were filtered through a 0.45-µm membrane, layered over an equal volume of 50% w/v sucrose solution and ultracentrifuged at 150,000 × *g* for 1 h and 4°C. Pellets containing the OMVs were washed once with 10 mM HEPES, 200 mM NaCl at pH 7.4, and stored at -80 °C until use.

OMVs were quantified by total protein dosage with the Pierce^®^ BCA Protein Assay Kit (Thermo Scientific). OMVs were then analyzed by SDS-PAGE and immunoblotting. To determine the levels of NDM-1, NDM-1 C26A, NDM-1 2RE, VIM-2, IMP-1 and IMP-1 4KE into OMVs, the protein band intensities in the whole cells (WC) and in the OMVs, from *E. coli* expressing each MBL, were quantified from PVDF membranes with the software ImageJ (28). In addition, the band intensity of the main outer membrane protein (OmpF/C) in the OMVs preparations was quantified from SDS-PAGE gels. The quantity of each MBL in the OMVs (from immunoblot) was normalized to the amount of OmpF/C (from SDS-PAGE), in the same sample, and then this value was divided by the quantity of each MBL in whole cells (from immunoblot). Finally, the values plotted in the Figures 1C, 2C and 3C correspond to the relativization to NDM-1 or IMP-1 value, as we show in the following equation:

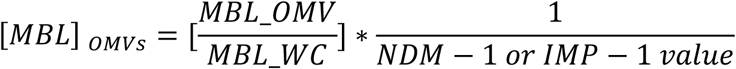

where [MBLs] _OMVs_ =Relative level of MBLs into OMVs, MBL_WC = Intensity of the MBL band in whole cells and MBL_OMV = Intensity of the MBL band in OMVs normalized to Omp band in the same sample.

### MBL detection and β-lactamase activity measurements

MBL protein levels were determined by SDS-PAGE followed by western blotting with Strep-tag II monoclonal antibodies (at a 1:1000 dilution from 200 μg/ml solution) (Novagen) and immunoglobulin G-alkaline phosphatase conjugates (at a 1:3000 dilution). Briefly, the samples were mixed with loading buffer and heated to denature the peptide structure. SDS-PAGE (14%) was used for separation of the sample components and subsequently transferred onto a PVDF membrane (GE). β-lactamase activity into OMVs purified from *E. coli* expressing the different MBLs and their variants was measured in a JASCO V-670 spectrophotometer at 30 °C in 10 mM HEPES, 200 mM NaCl at pH 7.4, in 0.1 cm cuvettes, using 400 μM imipenem as a substrate. Imipenem hydrolysis was monitored at 300 nm (Δε_300nm_ = −9000 M^−1^ cm^−1^).

### Coarse-grained molecular dynamics simulations

In CG MD simulations, the enzymes were located at a 40 Å distance from the bilayer, in order to avoid early protein/membrane encounters that might bias the subsequent interactions. The Martini 2.2p (polarizable) force field was used for all the simulations. The atomistic 3D structures of NDM-1, VIM-2 and IMP-1 were taken from the Protein Data Bank (PDB codes: 5ZGE (29) for NDM-1, 1KO3 (30) for VIM-2 and 5EV6 (31) for IMP-1) and turned into a coarse-grained model with the Martinize tool. All lipid bilayers were generated with the *Insane* tool of Martini (32) with a lipid composition that mimics the bacterial outer membrane (91% PEs, 6% CDLs and 3% PGs, in agreement with lipidomics analyses present in literature (33, 34). For what concerns the acyl chains, we used the most common motifs for each lipid type, that are: one 16:0 and one 18:1 (1-palmitoyl-2-oleoyl) for PEs and PGs, and four 18:1 for CDLs. All the systems were solvated with the polarizable Martini water model (35) and ionized with 150 mM of NaCl. Each enzyme was simulated in 5 distinct replicas. Each replica had a simulation time of 2 *μ*s. Frames for the analysis were collected every 750 ps. A membrane binding event is considered to have occurred when the protein settles at <3 Å distance from the membrane within the first *μ*s, and this distance remains constant for the rest of the simulation (i.e., no detachment occurs). For each system, the equilibration procedure was run as follows: firstly, the system went through 5000 steps of minimization using steepest descent; secondly, it was equilibrated with 5 ns of MD in the NVT ensemble, using Particle Mesh Ewald (PME) for the electrostatic contributions and velocity rescale algorithm for temperature coupling at 310 K. The production phase was conducted in the NPT ensemble, using a Parrinello-Rahman semi-isotropic coupling algorithm (36) to maintain the pressure constant at 1 bar. The two zinc ions in the catalytic site were represented as one single Martini bead of Qa type, connected to the 6 coordinating residues through harmonic potentials. Such particle was charged with +2e, that is the total net charge of the catalytic site.

## ACKNOWLEDGEMENTS

We thank Marina Avecilla (IBR) for excellent technical assistance. This work was supported by a MinCyT (SUIZ/17/10) and a Swiss National Science Foundation grant (514106) to A.J.V. and M.D.P., a grant from ANPCyT (PICT-2016-1657) to A.J.V. and an NIAID grant (2R01AI100560-06A1) to A.J.V. and R.A.B. C.L. is recipient of a postdoctoral fellowship from CONICET, and L.J.G. and A.J.V. are staff members from CONICET.

C.L., A.P., G.B., R.A.B., L.J.G., M.D.P. and A.J.V. conceived the project and designed the experiments. C.L. performed the construction of NDM-1 R2E and IMP-1 4KE mutants, MICs determinations, OMVs purification, MBL detection and determination of β-lactamase into OMVs. G.B. performed the liposome preparation and the liposome flotation assay. C.L. and L.J.G. helped to carry out the IMP-1 liposome flotation assay. A.P. performed the computational experiments. C.L and A.P. designed the figures. C.L., L.J.G. and A.J.V. wrote the manuscript. All authors discussed the results and commented on the manuscript. The content is solely the responsibility of the authors and does not necessarily represent the official views of the National Institutes of Health or the Department of Veterans Affairs.

---

**Supplementary Table 1.** MIC values of imipenem (IPM) and ceftazidime (CAZ) for *E. coli* carrying the empty vector (EV) or expressing *bla*_NDM-1_, *bla*_NDM-1 C26A_ (NDM-1 C26A), *bla*_NDM-1 R45E/R52E_ (NDM-1 2RE), *bla*_IMP-1_, *bla*_IMP-1 K87E/K89E/K145E/K147E_ (IMP-1 4KE) at 20 μM of IPTG. Data correspond to mean values from three independent experiments.

| MIC values (μg ml<sup>-1</sup>) of imipenem (IPM) and ceftazidime (CAZ) for *E. coli* expressing MBLs at 20 μM of IPTG | | |
| --- | --- | --- |
|  | IPM | CAZ |
| EV | <0.06 | 0.03 |
| NDM-1 | 2 | 1024 |
| NDM-1 C26A | 2 | 1024-2048 |
| NDM-1 2RE | 2 | 1024 |
| IMP-1 | 1 | 258 |
| IMP-1 4KE | 1 | 126 |
| VIM-2 | 1-2* | 16* |

\*at 10 μM of IPTG

**Figure S1.**
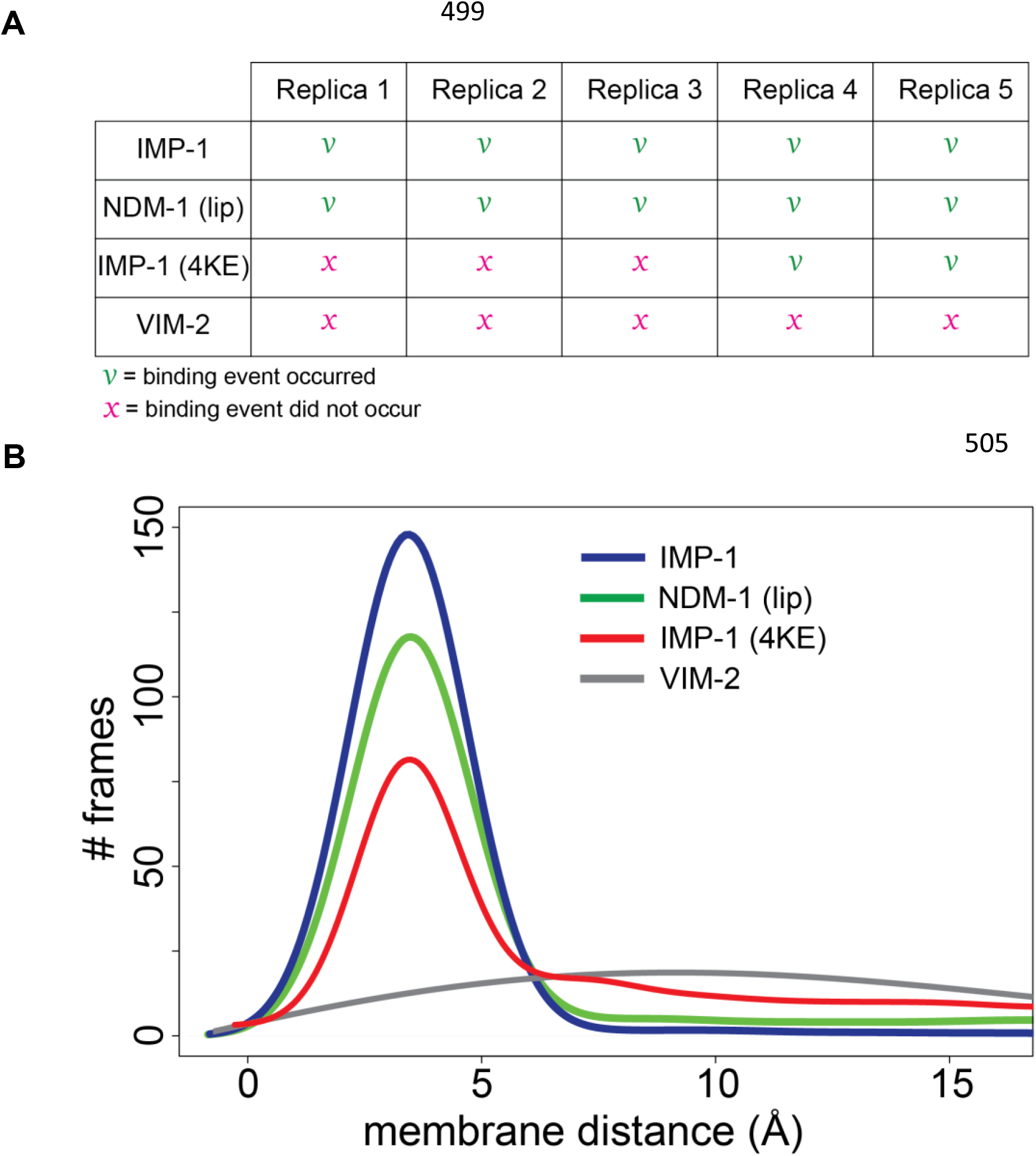
MBLs membrane association. (A) Binding events observed for WT IMP-1, NDM-1, mutated IMP-1 (IMP-1 4KE) and VIM-2 in the different CG MD replicas. (B) Distribution of CG MD trajectory frames collected every 750 ps for all available replicas with respect to protein-membrane distance.

**Figure S2.**
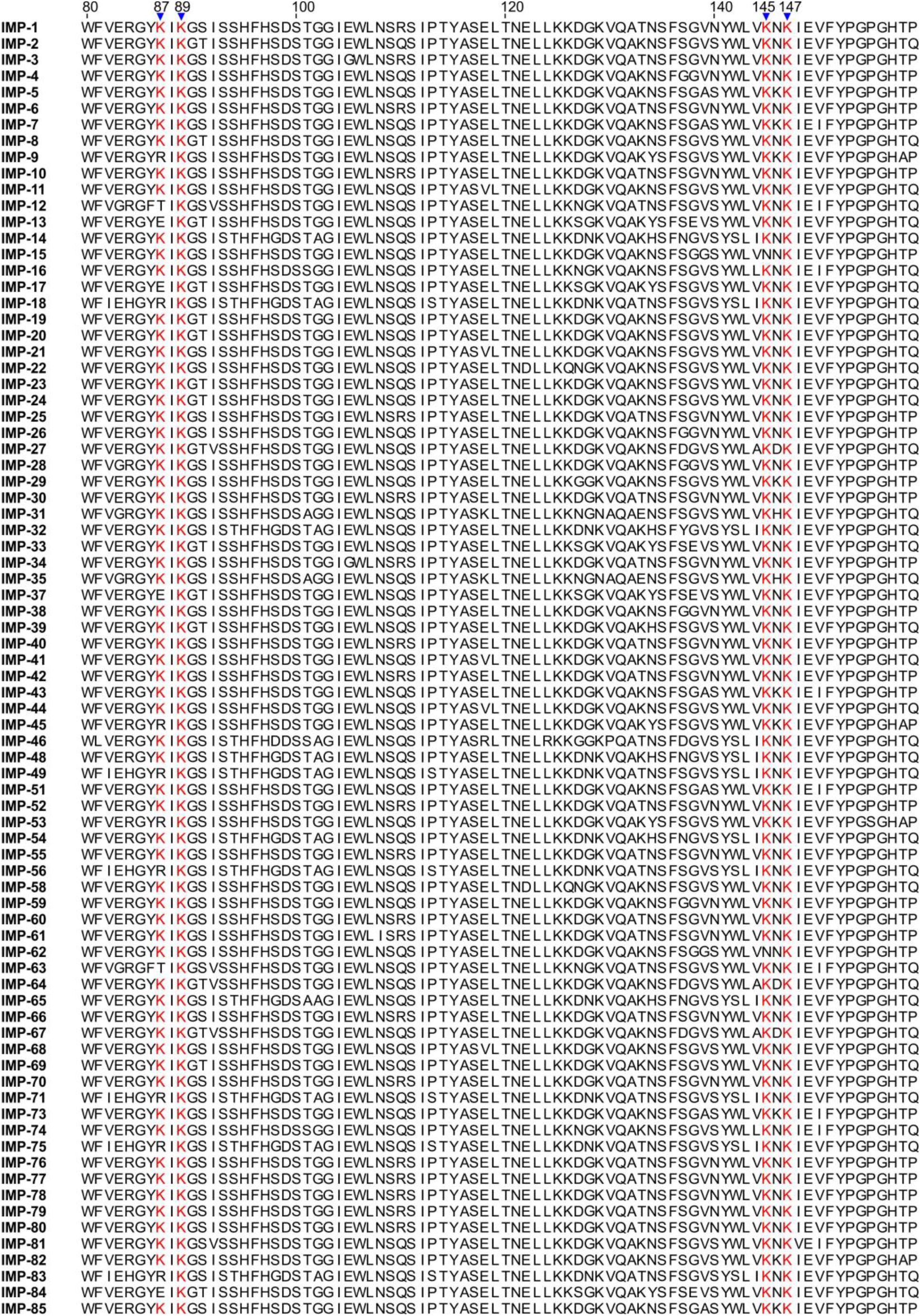
Sequence alignment of known 80 IMP allelic variants (only the region of interest is shown, from amino acid 80 to 159). Conserved Lys residues (K) at positions 87, 89, 145 and 147 are shown in red. Alignment was performed with the T-Coffee tool, available at www.tcoffee.org.

## REFERENCES

1. Papp-Wallace KM, Endimiani A, Taracila MA, Bonomo RA. 2011. Carbapenems: Past, present, and future. Antimicrob Agents Chemother 55:4943– 4960.

2. Walsh TR, Toleman MA, Poirel L, Nordmann P. 2005. Metallo-beta-lactamases: the quiet before the storm? Clin Microbiol Rev 18:306–25.

3. Meini MR, Llarrull LI, Vila AJ. 2015. Overcoming differences: The catalytic mechanism of metallo-β-lactamases. FEBS Lett 589:3419–3432.

4. Codjoe F, Donkor E. 2017. Carbapenem Resistance: A Review. Med Sci 6:1.

5. Ju LC, Cheng Z, Fast W, Bonomo RA, Crowder MW. 2018. The Continuing Challenge of Metallo-β-Lactamase Inhibition: Mechanism Matters. Trends Pharmacol Sci 39:635–647.

6. Palacios AR, Rossi MA, Mahler GS, Vila AJ. 2020. Metallo-β-lactamase inhibitors inspired on snapshots from the catalytic mechanism. Biomolecules 10:854.

7. Boyd SE, Livermore DM, Hooper DC, Hope WW. 2020. Metallo-β-lactamases: Structure, function, epidemiology, treatment options, and the development pipeline. Antimicrob Agents Chemother 64:e00397–20.

8. Nordmann P, Poirel L. 2014. The difficult-to-control spread of carbapenemase producers among Enterobacteriaceae worldwide. Clin Microbiol Infect 20:821– 830.

9. F. Mojica M, A. Bonomo> R, Fast W. 2015. B1-Metallo-β-Lactamases: Where Do We Stand? Curr Drug Targets 17:1029–1050.

10. Halat DH, Moubareck CA. 2020. The current burden of carbapenemases: Review of significant properties and dissemination among gram-negative bacteria. Antibiotics 9:186.

11. King D, Strynadka N. 2011. Crystal structure of New Delhi metallo-β-lactamase reveals molecular basis for antibiotic resistance. Protein Sci 20:1484–1491.

12. González LJ, Bahr G, Nakashige TG, Nolan EM, Bonomo RA, Vila AJ. 2016. Membrane anchoring stabilizes and favors secretion of New Delhi metallo-β-lactamase. Nat Chem Biol 12:516–522.

13. Schwechheimer C, Kuehn MJ. 2015. Outer-membrane vesicles from Gram-negative bacteria: biogenesis and functions. Nat Rev Microbiol 13:605–619.

14. Bonnington KE, Kuehn MJ. 2014. Protein selection and export via outer membrane vesicles. Biochim Biophys Acta - Mol Cell Res 1843:1612–1619.

15. Caruana JC, Walper SA. 2020. Bacterial Membrane Vesicles as Mediators of Microbe – Microbe and Microbe – Host Community Interactions. Front Microbiol 11:432.

16. Ciofu O. 2000. Chromosomal beta-lactamase is packaged into membrane vesicles and secreted from Pseudomonas aeruginosa. J Antimicrob Chemother 45:9–13.

17. Li Z, Clarke AJ, Beveridge TJ. 1998. Gram-negative bacteria produce membrane vesicles which are capable of killing other bacteria. J Bacteriol 180:5478–5483.

18. Liao YT, Kuo SC, Chiang MH, Lee YT, Sung WC, Chen YH, Chen TL, Fung CP. 2015. Acinetobacter baumannii extracellular OXA-58 is primarily and selectively released via outer membrane vesicles after sec-dependent periplasmic translocation. Antimicrob Agents Chemother 59:7346–7354.

19. Rumbo C, Fernández-Moreira E, Merino M, Poza M, Mendez JA, Soares NC, Mosquera A, Chaves F, Bou G. 2011. Horizontal transfer of the OXA-24 carbapenemase gene via outer membrane vesicles: A new mechanism of dissemination of carbapenem resistance genes in Acinetobacter baumannii. Antimicrob Agents Chemother 55:3084–3090.

20. Chatterjee S, Mondal A, Mitra S, Basu S. 2017. Acinetobacter baumannii transfers the blaNDM-1gene via outer membrane vesicles. J Antimicrob Chemother 72:2201–2207.

21. Schaar V, Nordström T, Mörgelin M, Riesbeck K. 2011. Moraxella catarrhalis outer membrane vesicles carry β-lactamase and promote survival of Streptococcus pneumoniae and Haemophilus influenzae by inactivating amoxicillin. Antimicrob Agents Chemother 55:3845–3853.

22. López C, Ayala JA, Bonomo RA, González LJ, Vila AJ. 2019. Protein determinants of dissemination and host specificity of metallo-β-lactamases. Nat Commun 10:3617.

23. Prunotto A, Bahr G, González LJ, Vila AJ, Dal Peraro M. 2020. Molecular Bases of the Membrane Association Mechanism Potentiating Antibiotic Resistance by New Delhi Metallo-β-lactamase 1. ACS Infect Dis 6:2719–2731.

24. Naas T, Oueslati S, Bonnin RA, Dabos ML, Zavala A, Dortet L, Retailleau P, Iorga BI. 2017. Beta-lactamase database (BLDB)–structure and function. J Enzyme Inhib Med Chem 32:917–919.

25. Devos S, Stremersch S, Raemdonck K, Braeckmans K, Devreese B. 2016. Intra- and interspecies effects of outer membrane vesicles from Stenotrophomonas maltophilia on beta-lactam resistance. Antimicrob Agents Chemother 60:2516– 2518.

26. Valguarnera E, Scott NE, Azimzadeh P, Feldman MF. 2018. Surface Exposure and Packing of Lipoproteins into Outer Membrane Vesicles Are Coupled Processes in Bacteroides. mSphere 3:e00559–18.

27. CLSI. 2015. M07-A10: Methods for dilution antimicrobial susceptibility tests for bacteria that grow aerobically.

28. Schneider CA, Rasband WS, Eliceiri KW. 2012. NIH Image to ImageJ: 25 years of image analysis. Nat Methods 9:671–5.

29. Zhang H, Ma G, Zhu Y, Zeng L, Ahmad A, Wang C, Pang B, Fang H, Zhao L, Hao Q. 2018. Active-Site Conformational Fluctuations Promote the Enzymatic Activity of NDM-1. Antimicrob Agents Chemother 62:e01579–18.

30. Garcia-Saez I, Docquier JD, Rossolini GM, Dideberg O. 2008. The Three-Dimensional Structure of VIM-2, a Zn-β-Lactamase from Pseudomonas aeruginosa in Its Reduced and Oxidised Form. J Mol Biol 375:604–611.

31. Hinchliffe P, González MM, Mojica MF, González JM, Castillo V, Saiz C, Kosmopoulou M, Tooke CL, Llarrull LI, Mahler G, Bonomo RA, Vila AJ, Spencer J. 2016. Cross-class metallo-β-lactamase inhibition by bisthiazolidines reveals multiple binding modes. Proc Natl Acad Sci 113:E3745–E3754.

32. Wassenaar TA, Ingólfsson HI, Böckmann RA, Tieleman DP, Marrink SJ. 2015. Computational lipidomics with insane: A versatile tool for generating custom membranes for molecular simulations. J Chem Theory Comput 11:2144–2155.

33. Lugtenberg EJJ, Peters R. 1976. Distribution of lipids in cytoplasmic and outer membranes of Escherichia coli K12. Biochim Biophys Acta (BBA)/Lipids Lipid Metab 441:38–47.

34. Morein S, Andersson AS, Rilfors L, Lindblom G. 1996. Wild-type Escherichia coli cells regulate the membrane lipid composition in a “window” between gel and non-lamellar structures. J Biol Chem 271:6801–6809.

35. Yesylevskyy SO, Schäfer L V., Sengupta D, Marrink SJ. 2010. Polarizable Water Model for the Coarse-Grained MARTINI Force Field. PLoS Comput Biol 6:e1000810.

36. Parrinello M, Rahman A. 1981. Polymorphic transitions in single crystals: A new molecular dynamics method. J Appl Phys 52:7182–7190.

